# Stratified Linkage Disequilibrium Score Regression reveals enrichment of eQTL effects on complex traits is not tissue specific

**DOI:** 10.1101/107482

**Authors:** Hill F. Ip, Rick Jansen, Abdel Abdellaoui, Meike Bartels, UK Brain Expression Consortium, Dorret I. Boomsma, Michel G. Nivard

## Abstract

Both gene expression levels and eQTLs (expression quantitative trait loci) are partially tissue-specific, complicating the detection of eQTLs in tissues with limited sample availability, such as the brain. However, eQTL overlap between tissues might be non-trivial, allowing for inference of eQTL functioning in the brain via eQTLs measured in readily accessible tissues, e.g. whole blood. Using Stratified Linkage Disequilibrium Score Regression (SLDSR), we quantify the enrichment in GWAS signal of blood and brain eQTLs in genome-wide association study (GWAS) on 11 complex traits (schizophrenia, BMI, educational attainment, Crohn’s disease, rheumatoid arthritis, ulcerative colitis, age at menarche, coronary artery disease, height, LDL levels, and smoking behavior). Our analyses established significant enrichment of blood and brain eQTLs in their effects across all traits. As we do not know the true number of causal eQTLs, it is difficult to determine the precise magnitude of enrichment. We found no evidence for tissue-specific enrichment in GWAS signal for either eQTLs uniquely seen in the brain or whole blood. To follow up on our findings, we tested tissue-specific enrichment of eQTLs discovered in 44 tissues by the Genotype-Tissue Expression (GTEx) consortium, and, again, found no tissue-specific eQTL effects. We further integrated the GTEx eQTLs with SNPs associated with tissue-specific histone modifiers, and interrogate its effect on rheumatoid arthritis and schizophrenia. We observed substantially enriched effects on schizophrenia, though again not tissue-specific. Finally, we extracted eQTLs in tissue-specific differentially expressed genes, and determined their effects on rheumatoid arthritis and schizophrenia. We conclude that, while eQTLs are strongly enriched in GWAS signal, the enrichment is not specific to the tissue used in eQTL discovery. Therefore, working with relatively accessible tissues, such as whole blood, as proxy for eQTL discovery is sensible; and restricting lookups for GWAS hits to a specific tissue might not be advisable.

## Introduction

The aim of genome-wide association studies (GWASs) is to detect statistically significant associations between genetic variants, such as single nucleotide polymorphisms (SNPs), and a trait of interest (Hirschhorn and Daly 2005). GWASs have identified many genetic variants and thereby provided insights into the genetic architecture of complex traits (Hirschhorn and Daly 2005; Visscher et al. 2012). However, as a large number of variants identified through GWASs are located outside of coding regions and specific knowledge of regulatory elements is limited, uncovering a relationship between GWAS hits and biological function has proven to be complicated (Lowe and Reddy 2015). Expression quantitative trait loci (eQTLs) contain SNPs that influence gene expression, and are not necessarily located in coding regions. eQTLs may aid functional annotation of SNPs that have been identified in a GWAS and are located outside of coding regions (Morley et al. 2004; Lowe and Reddy 2015). Previous work has found substantial enrichment of eQTLs among GWAS hits (Manolio et al. 2009; Nicolae et al. 2010; Torres et al. 2014) and an enrichment in their genome-wide effect on complex traits (Davis et al. 2013). Therefore, eQTLs are viewed as an important tool in moving from genome-wide association to biological interpretation.

As a result of difference in gene expression between cells originating from different tissues, eQTLs are potentially tissue-specific (Hernandez et al. 2012; GTEx Consortium 2015). Tissue-specificity poses no problem if the tissue of interest is readily available for research, such as whole blood. However, discovery of eQTLs gets complicated when measurement of expression levels in a tissue is limited by ethical and practical considerations, for example in brain tissue. Several studies have shown that the overlap between eQTLs from different tissues might actually be larger than initially assumed (Ding et al. 2010; Nica et al. 2011). The Genotype-Tissue Expression (GTEx) consortium identified eQTLs in a wide range of human tissues and showed that 54–90% of the eQTLs identified in one tissue are also designated as an eQTL in at least one other tissue (GTEx Consortium 2015; Aguet et al. 2016). In another study, Liu *et al* (2016) found a high average pairwise genetic correlation (r_g_=0.738) of local gene expression between tissues. Nevertheless, small differences in terms of eQTL effect may be of considerable importance in terms of the effect an eQTL might have on complex traits related to specific tissues. It is, therefore, worthwhile to investigate the specific utility of tissue-specific eQTLs in their effect on complex traits, as studied in GWAS, as the discovery of eQTLs for tissues such as the brain might be advanced by eQTLs discovered in more accessible tissues, such as whole blood. The use of accessible tissues, though, depends on a substantial degree of similarity of eQTL effect across tissue, and to what extend eQTL differences between tissues are important in complex trait etiology.

Stratified Linkage Disequilibrium Score Regression (SLDSR) is a technique that estimates the SNP-heritability (h^2^_SNP_) of a trait based on GWAS summary statistics (Bulik-Sullivan et al. 2015; Finucane et al. 2015). By simultaneously analyzing multiple categories of SNPs (annotations), SLDSR can partition h^2^_SNP_ by annotation (h^2^_annot_) and thereby provides a way to jointly quantify the enrichment in GWAS signal of several annotations. Here, we extend SLDSR by including annotations containing *cis*-eQTLs, i.e. eQTLs located closely to the gene with which they associate (Brem et al. 2002; Ramasamy et al. 2014), discovered in multiple tissues. To this end, we perform analyses based on representative eQTL resources, and consider a variety of traits as outcomes.

Firstly, we selected all eQTLs per gene discovered in large samples of RNA expression levels assessed in whole blood (N=4896)(Wright et al. 2014; Jansen et al. 2017) and in brain tissues (N=134) (Ramasamy et al. 2014), and quantified the contribution of these blood and brain eQTLs to the genetic variance in complex traits captured in GWAS. We then estimated tissue-specific eQTL effects on complex traits by quantifying the enrichments of eQTLs uniquely found in whole blood or uniquely found in brain, conditional on the enrichment of the complete blood eQTL annotation or complete brain eQTL annotation, respectively. We considered the effect of eQTLs on 11 complex traits: schizophrenia, BMI, educational attainment, Crohn’s disease, rheumatoid arthritis, ulcerative colitis, age at menarche, coronary artery disease, height, LDL levels, and smoking behavior.

Secondly, we retrieved all eQTLs identified in any of the 44 tissues from the GTEx consortium (N=70–361, median=126.5)(GTEx Consortium 2015; Aguet et al. 2016). We considered the enrichment in GWAS signal of the union of all GTEx eQTLs, and, additionally, the enrichment of tissue-specific eQTL effects on top of the union of all GTEx eQTLs. We expected to observe tissue-specific enrichment of eQTLs in their effects on complex traits related to the tissue in question, e.g. eQTLs discovered in immune-related tissues are expected to show higher enrichments in their effect on immune-related traits compared to eQTLs found in skin tissue. We considered tissue-specific enrichment of *cis*-eQTLs in their effect on schizophrenia (a disorder where there is strong prior evidence for the involvement of processes in the brain) and rheumatoid arthritis (a disease with strong prior evidence for the involvement of processes in immune tissue) as GWAS for these traits are well powered for extended LD-score-based analyses. We further considered the enrichment of the intersection of *cis*-eQTLs discovered in any tissue, and histone modification in a specific tissue (i.e. tissue-specific epigenetically changed chromatin states in regulatory regions). Finally, we explored the enrichment in GWAS signal of eQTLs for tissue-specific differentially expressed genes.

Our analyses were designed to elucidate the relation between eQTLs and complex traits, and to quantify the extent to which this relation is dependent on the tissue used in eQTL discovery. Our analysis further considered the enrichment of genomic regions related to gene expression and epigenetically modified in specific tissues.

## Material and Methods

### SLDSR method

A measure of linkage disequilibrium (LD) for each SNP, called an “LD score”, can be computed by taking the sum of correlations between that SNP and all neighboring SNPs (Bulik-Sullivan et al. 2015;

Finucane et al. 2015). Under a polygenic model, LD scores are expected to show a linear relationship with GWAS test statistics of corresponding SNPs, where the slope is proportional to h^2^_SNP_. For SLDSR, LD scores are based on only (functional) parts of the genome and used as predictors in a multiple linear regression (Finucane et al. 2015). In this manner, SLDSR is able to partition h^2^_SNP_ into parts that are explained by these parts of the genome (i.e. h^2^_annot_), while accounting for influences of the remaining annotations in the model. The enrichment of an annotation is then obtained by taking the ratio of h^2^_annot_ over the proportion of SNPs that fall within that annotation. For eQTLs, the denominator, i.e. the number of SNPs in the annotation, is a complicated quantity: not all significant eQTLs are likely causal; whereas including only lead, or putative causal, eQTLs may result in very small annotations located near genes and other regulatory elements, which presents a risk of inflated estimates of the enrichment in GWAS signal. What constitutes an eQTL is sufficiently vague and open for interpretation for us to consider the effect of multiple inclusion rules for inclusion of a SNP into the eQTL annotation. Since eQTLs are essentially discovered in what amounts to a local GWAS, we expect the average LD score of eQTLs to be higher than that of an average SNP, which may influence the results of downstream SLDSR analysis. In order to break the relation between LD score and probability of inclusion, we consider eQTL annotations which are based on a sample of from all significant eQTLs for a given probe. First, we included the most strongly associated SNP, a SNP with a high expected LD score. Second, we included one SNP per probe with a median *p*-value. Third, we included one SNP per probe with a mean *p*-value. Fourth, we included the 10 most strongly associated SNPs per probe. Finally, we included all SNPs significantly associated with gene expression after FDR correction at α=0.05. We add each annotation separately to the baseline categories in an SLDSR model, and determined how the various *p*-value thresholds influence the SLDSR coefficient and its test statistic. For each annotation, we looked up the SNPs in the baseline category, and extracted their baseline LD scores and minor allele frequencies (MAF). We then compared the mean LD score, median LD score and mean MAF between the annotations and the entire baseline category. Based on the results (S1 Figure - S2 Figure, S3 Table), we consider all significant *cis*-eQTLs as an annotation, and retain additional gene-centric and regulatory annotations ins the model.

### Target traits

As outcome for SLDSR, we used summary statistics of GWASs on Crohn’s disease (Jostins et al. 2012), rheumatoid arthritis (Okada et al. 2014), ulcerative colitis (Jostins et al. 2012), BMI (EK et al. 2010), educational attainment (Rietveld et al. 2013), schizophrenia (Ripke et al. 2014), age at menarche (Perry et al. 2014), coronary artery disease (Schunkert et al. 2011), height (Allen et al. 2010), LDL levels (Teslovich et al. 2010), and smoking behavior (Furberg et al. 2010). The first three traits were chosen because they are related to the immune system and are therefore expected to reveal considerable enrichment of blood eQTL signal (Jostins et al. 2012; Okada et al. 2014). Similarly, brain eQTLs are expected to show substantial enriched effects due to previous reports on the involvement of the central nervous system (CNS) in schizophrenia (Ripke et al. 2014), educational attainment (Rietveld et al. 2013), and BMI (Vimaleswaran et al. 2012). Of course, these traits do not perfectly align with either tissue, e.g. the immune system has been implicated in the etiology of schizophrenia (Andreassen et al. 2015) and BMI (Karalis et al. 2009), and might therefore also be enriched in their effects for the other eQTL set. Enrichment of blood and brain eQTL effects on the remaining traits was calculated to contrast the results with traits for which we do not have a strong *a priori* expectation of the relationship between trait and tissue.

The discovery sample for detection of blood eQTLs (Wright et al. 2014; Jansen et al. 2017) included participants from the Netherlands Twin Register (NTR)(Boomsma et al. 2008) and participants from the Netherlands Study of Depression and Anxiety (NESDA)(Penninx et al. 2008). These two cohorts did not participate in the GWAS for schizophrenia, Crohn’s disease, rheumatoid arthritis, ulcerative colitis, or coronary artery disease. However, participants from these two cohorts, not necessarily the same ones, did participate in the GWAS for height, BMI, LDL levels, smoking behavior, educational attainment, and age at menarche. For educational attainment and smoking behavior, we were able to obtain summary statistics omitting subjects from NTR/NESDA. For both these traits, we looked at trait-specific enrichment of blood and brain eQTL effect in GWAS signal, comparing results from using publicly available datasets with using summary statistics based on the same sample without subjects from the NTR or NESDA. The results did not reveal appreciable differences between the respective datasets for educational attainment, but did show substantial differences for smoking behavior (S4 Figure). This latter finding could conceivably be a function of relatively strong effects of smoking behavior on gene-expression levels (Vink et al. 2015). Therefore, the remaining analyses for smoking behavior were performed using the summary statistics omitting the NTR and NESDA, whereas analyses for the remaining traits (height, BMI, LDL levels, and educational attainment) were run using publicly available summary statistics. This caveat only applies to eQTL annotations based on NTR/NESDA data (i.e. whole blood). We note that the issue of overlap also applies to other techniques where the error covariance is assumed to be zero (e.g. TWAS, mendelian randomization analysis, SMR, etc.).

### Blood and brain eQTL enrichment

A catalog of whole blood *cis*-eQTLs was obtained from Jansen *et al* (2017; Wright et al. 2014), where all eQTLs significantly associated with gene expression in whole blood for each probe set were selected for inclusion in our whole blood eQTL annotation. A list of brain eQTLs was obtained from the UK brain expression consortium (UKBEC), for which the analyses are described in Ramasamy *et al* (2014). We based the brain eQTL annotation on SNPs that were significantly associated with the average gene expression across 12 brain regions. SLDSR annotations were constructed as per the instructions in Bulik-Sullivan et al. and Finucane et al. (2015). To guard against upward bias in the eQTL enrichment signal, two extra annotations containing SNPs within a 500 base pair (bp) and 100bp window around any eQTL were constructed for each eQTL set (Finucane et al. 2015). To ensure that the enrichment of eQTL effects in GWAS signal was not in fact caused by their proximity to the genes they influence, an additional gene centric annotation was computed, which contained all genes for which eQTLs were included. Finally, we performed an inverse-variance weighted meta-analysis across the traits to determine the average effect of blood and brain eQTLs on complex traits in general.

### Tissue-specific eQTL enrichment

To distinguish between the effects of blood- and brain-specific eQTLs, we split each annotation into two sets based on the overlap in genes that were tagged by eQTLs from both tissue. That is, the brain eQTL annotation was split into an annotation of brain eQTLs which regulate genes for which also at least one blood eQTL was found, and a second annotation of eQTLs that tagged genes for which only brain eQTLs were found. Likewise, the blood eQTL annotation was split into an annotation containing only eQTLs that tagged genes for which eQTLs from both tissue was found, and an annotation consisting of blood eQTLs that tagged genes for which only eQTLs have been found in blood. In doing so, we are saying that the effect of an eQTL is mediated via the gene it is associated with. Then, if two different SNPs are associated with the expression of the same gene, but in different tissues, this gene is likely the mechanism by which the SNP influences a trait. Contrary, when the same SNP affects different genes in different tissues, this SNP can be seen to have a tissue-specific mediation of its effect. We thus define eQTLs shared across tissue as eQTLs that influence a common gene in separate tissues.

### Enrichment of eQTLs from 44 tissues

There are several limitations to above mentioned analyses of tissue-specific enrichments of eQTL effects in GWAS signal. The eQTLs are obtained from two different projects, which vary in terms of sample size and their definition of an eQTL. To mitigate the heterogeneity between studies, and to extend to additional tissues. We performed additional analyses using eQTLs obtained by a common pipeline from 44 tissues (see S5 Table) and based on a broader eQTL locus definition (GTEx Consortium 2015; Aguet et al. 2016). For each of the 44 tissues, we created annotations for analysis in SLDSR following the previously described procedure. Analogous to the procedure of Finucane *et al* (2015) for cell-type-specific analysis using SLDSR, we did not specify windows for the individual GTEx annotations, but included an additional annotation that contained the union of all GTEx eQTLs, i.e. all SNPs that are designated as part of at least one of the 44 individual GTEx annotations, and added a 100bp and 500bp window around this union of GTEx eQTLs. SLDSR is essentially a multiple regression and, due to the high number of models and high number of predictors (i.e. annotations) in each model, has a high multiple testing burden. As such, it requires well-powered GWAS in order to accurately partition the heritability the various annotations (Bulik-Sullivan et al. 2015; Finucane et al. 2015). Based on the Z-score of the SNP-heritability and previous reports of substantial influence of either tissue in the etiology of the traits (Okada et al. 2014; Ripke et al. 2014; Finucane et al. 2015; Finucane et al. 2017), we obtained two well-powered traits, one for which we assume there to be significant enrichment in signal for blood eQTLs (rheumatoid arthritis) and one for brain eQTLs (schizophrenia). For each of these two traits, we ran one SLDSR model containing only the baseline categories and the union of GTEx eQTLs, and 44 additional models with the two previous annotations and one of the individual GTEx annotations at a time.

GTEx has relative small sample sizes for the brain eQTL discovery (mean=89 sample size, range=72–103) compared to other tissues (mean=160 sample size, range=70–361) (GTEx Consortium 2015; Aguet et al. 2016). To investigate the effect of differences in sample size on estimates of enrichments in GWAS signal, we collapsed the union of individual brain eQTL annotations into a shared brain eQTL annotation (i.e. an eQTL found in at least one of the GTEx brain annotations was included in the shared brain eQTL annotation). This annotation was then analyzed as an additional GTEx eQTL annotation. We further tested the relationship between tissue sample size and tissue eQTL enrichment.

### Enrichment of the intersection between eQTLs and histone marks

The availability of annotations based on tissue-specific histone marks made it possible to create an annotation that represents the intersection between eQTLs and this type of epigenetic modification related to enhancers and promoters of actively transcribed genes. We obtained LD score annotations of SNPs in regions that bear histone marks in cells from the CNS or immune system from Finucane *et al* (2015). Out of the 220 cell-type-specific histone mark that were available, 101 were found in the CNS or immune tissues. For each of the 101 annotations of SNPs in cell-type-specific histone marks, we extracted its intersection with the union of GTEx eQTLs and made a new annotation of eQTLs which intersected with histone marks (i.e. SNPs found in both annotations). We then analyzed each of the intersection annotations individually in a model together with the baseline categories, the union of GTEx eQTLs, and the corresponding cell-type-specific histone marks. Enrichments in GWAS signal of the intersection should be interpreted as enrichment of genome-wide SNP effects on a complex trait beyond the additive effects which work on all SNPs that are a *cis*-eQTL and histone mark in question. In fact, we test whether the interaction between tissue-specific chromatin state and eQTLs are enriched in their genome-wide effect on complex traits.

### Intersection of GTEx eQTLs and tissue-specific differentially expressed genes

Finucane *et al (* 2017) looked at tissue-specific gene expression and determined that the top 10% of these differentially expressed genes are substantially enriched in their effects in GWAS signals for a wide range of traits. Here, we build on these findings by taking the top 10% most highly differentially expressed genes in the 44 GTEx tissues; obtaining the eQTLs for these specific genes, regardless of the discovery tissue; and added these separately as an annotation to an SLDSR model together with the baseline categories and union of all GTEx eQTLs. A significant increase in enrichment in GWAS signal in the eQTLs compared to the genes themselves, would indicate that eQTLs explain part of the enrichment seen by Finucane *et al.*

## Results

### Effect of eQTL selection on regression parameter and test statistics

We compare five annotations that included various SNPs based on the *p*-value of their associations with gene-expression levels (lead eQTL, median eQTL, mean eQTL, top 10 lead eQTLs, and all eQTLs). Supplementary table S3 shows various metrics of these annotations. Surprisingly, lead eQTLs had the lowest mean and median LD score amongst the annotations, indicating that the annotation contains less signal (S3 Table). However, it was still higher compared to the mean or median of all SNPs in the baseline annotation. Including all significant eQTLs in the annotation resulted in the highest mean and median LD score. All annotations had a mean MAF 0.27-0.28, whereas the mean MAF of the entire baseline category was 0.24. Smaller annotations had a higher enrichment in GWAS signal (S1 Figure). However, the enrichment in GWAS signal did not differ between taking the lead eQTL, eQTLs with a mean *p*-value, or eQTLs with a median *p*-value. Finally, coefficient test statistics did not differ much between the annotations (S2 Figure). Based on the explorative analysis, we selected the annotation based on all significant eQTLs for further analysis.

### Blood and brain eQTL enrichment

We fitted an SLDSR model containing the baseline categories; the complete annotation for both brain and blood eQTL tissues, their 100 and 500bp windows, and gene-centric annotations to all traits (Crohn’s disease, rheumatoid arthritis, ulcerative colitis, BMI, educational attainment, schizophrenia, age at menarche, coronary artery disease, height, LDL levels, and smoking behavior). We found significant effects of brain eQTLs on educational attainment, rheumatoid arthritis, smoking behavior, and schizophrenia, and significant effect of blood eQTLs on height and smoking behavior (see S6 Table). We then meta-analyzed the results for all annotations, both in the baseline model, and those associated with eQTLs across the 11 traits. Our analyses revealed significant effect of both blood (*p* < 0.001) and brain (*p* < 0.001) eQTL effects on all traits (Figure 1, S7 Table), exceeding, in terms of significance, all the baseline categories considered by Finucane *et al* (2015) but conserved genomic regions. The gene-centric annotation for both blood and brain eQTLs showed no effect on any trait.

**Figure. 1.**
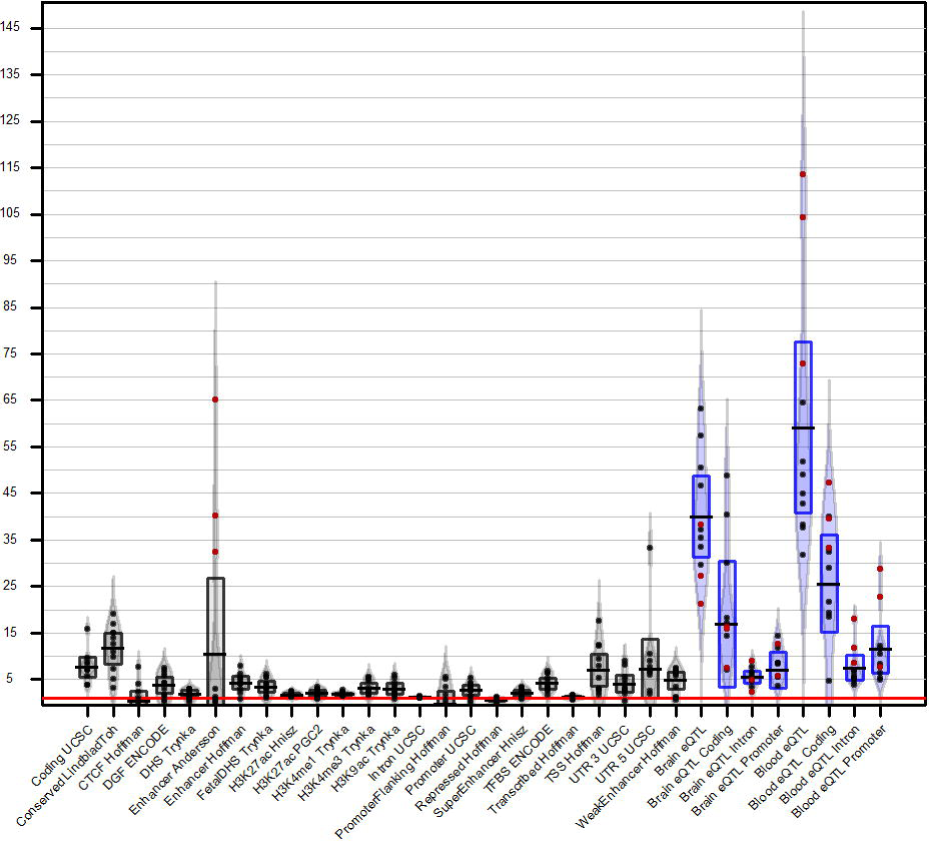
Average enrichment in GWAS signal of the 24 baseline annotations, 4 brain eQTL annotations and 4 blood eQTL annotations. Bar plot of the average enrichment in GWAS signal across all traits for the 24 main baseline annotations and 8 main eQTL annotations. Grey beans represent the baseline categories. Blue beans represent eQTLs. Black bars indicate average enrichment. Boxes show upper- and lower-bounds of the 95% confidence interval of the mean. Red dots show enrichments for immune-related traits. Horizontal red line indicates enrichment of 1, i.e. no enrichment.

We then separated the list of blood eQTLs into a list of unique and shared blood eQTLs based on the overlap in target genes with brain eQTLs, and modelled the baseline categories together with all blood eQTLs and the unique blood eQTLs. We did the same for brain eQTLs. We observed no evidence for depletion of blood-specific eQTLs (relative to all blood eQTLs) on brain-related traits, nor do we find significant depletion of effect on immune-related traits of eQTLs associated with genes for which eQTLs were solely identified in brain tissue (Table I and Table II).

### Enrichment of eQTLs from 44 tissues in GTEx

We interrogated the enrichment of the union of GTEx eQTLs and 44 individual GTEx annotations in their effect on schizophrenia and rheumatoid arthritis. Figure 2 shows the coefficient of the 45 GTEx annotations, sorted on their Z-scores for rheumatoid arthritis. In both cases, the union of GTEx eQTLs contributed significantly to explaining the polygenic signal(S8 Table), indicating that eQTLs were significantly enriched in their effects on complex traits. The individual annotations, however, performed notably worse and in some cases even suggested depletion of genome-wide effects of tissue-specific eQTLs on schizophrenia and rheumatoid arthritis. For rheumatoid arthritis, the coefficient Z-scores of the whole blood annotation reached nominal significance (Z=2.251), but failed correction for multiple testing. None of the other annotations reached nominal significance. The union of all GTEx brain annotations did not contribute significantly to explaining h^2^_SNP_ (Z=0.147, p=0.441). Sample size in the eQTL discovery phase appears to be a strong determinant of tissue-specific enrichment in GWAS signal. The correlation coefficients between the coefficient Z-scores and sample sizes were 0.6453 (p=2.253*10^−6^) and 0.4247 (p=0.004) for schizophrenia and rheumatoid arthritis, respectively.

**Figure. 2.**
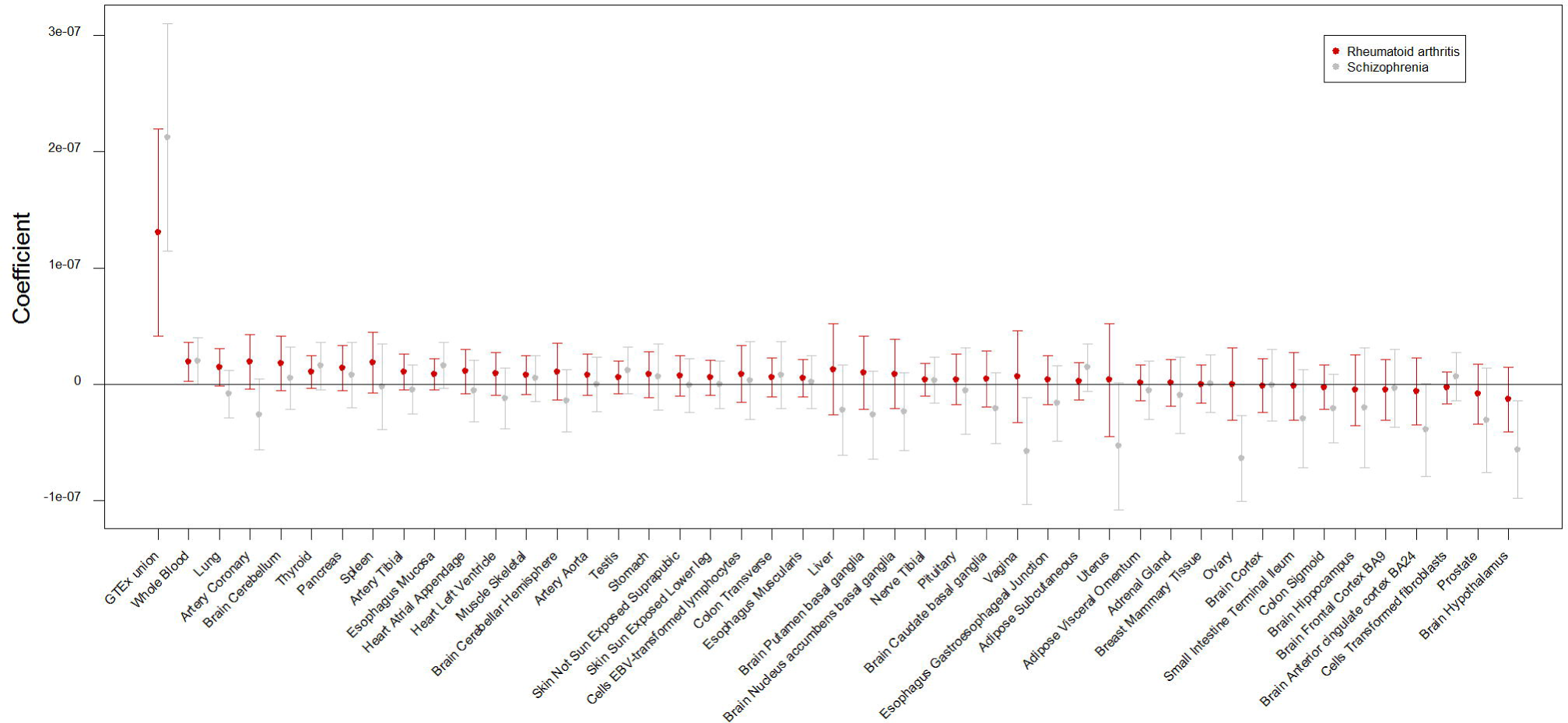
Coefficient Z-scores of the 45 GTEx annotations. Barplot of coefficient z-scores for all GTEx annotations for schizophrenia (grey) and rheumatoid arthritis (red). Bars are sorted from highest to lowest based on the results from schizophrenia. Horizontal dotted line indicates Bonferroni threshold for 45 tests. Two asterisks indicate bars passing Bonferroni correction for multiple testing.

### Enrichment of the intersection between eQTLs and histone marks

We interrogated the intersection of eQTLs and histone marks found in specific CNS and immune cells, and estimated the enrichment of the intersection in its effect on rheumatoid arthritis and schizophrenia. We found significant enrichment in GWAS signal for eQTLs that intersect with histones that bear modification H3K4me1, a modification thought to be present in the enhancer of actively transcribed genes (Zhou et al. 2011; Allis and Jenuwein 2016), in CNS cells for schizophrenia (see Figure 3). There was some evidence for significant enrichment of eQTLs that intersected with genomic regions in immune cells baring the H3K4me1 mark in their effect on schizophrenia, but not on rheumatoid arthritis. Specifically, none of the intersecting annotations showed evidence of enrichment for rheumatoid arthritis. For the separate annotations, we found significant enrichment in GWAS signal across all histone marks found in CNS cells and three significant immune cell-types that bear the H3K4me3 modification, a modification associated with transcriptional start sites and promoters of actively transcribed genes (Zhou et al. 2011; Allis and Jenuwein 2016), for schizophrenia (S9 Figure). The opposite picture was seen for rheumatoid arthritis: a wide variety of immune-cell specific histone marks showed significant enrichments in GWAS signal, while all marks found in CNS cells were below zero. The union of GTEx eQTLs reached statistical significance for all models.

**Figure. 3.**
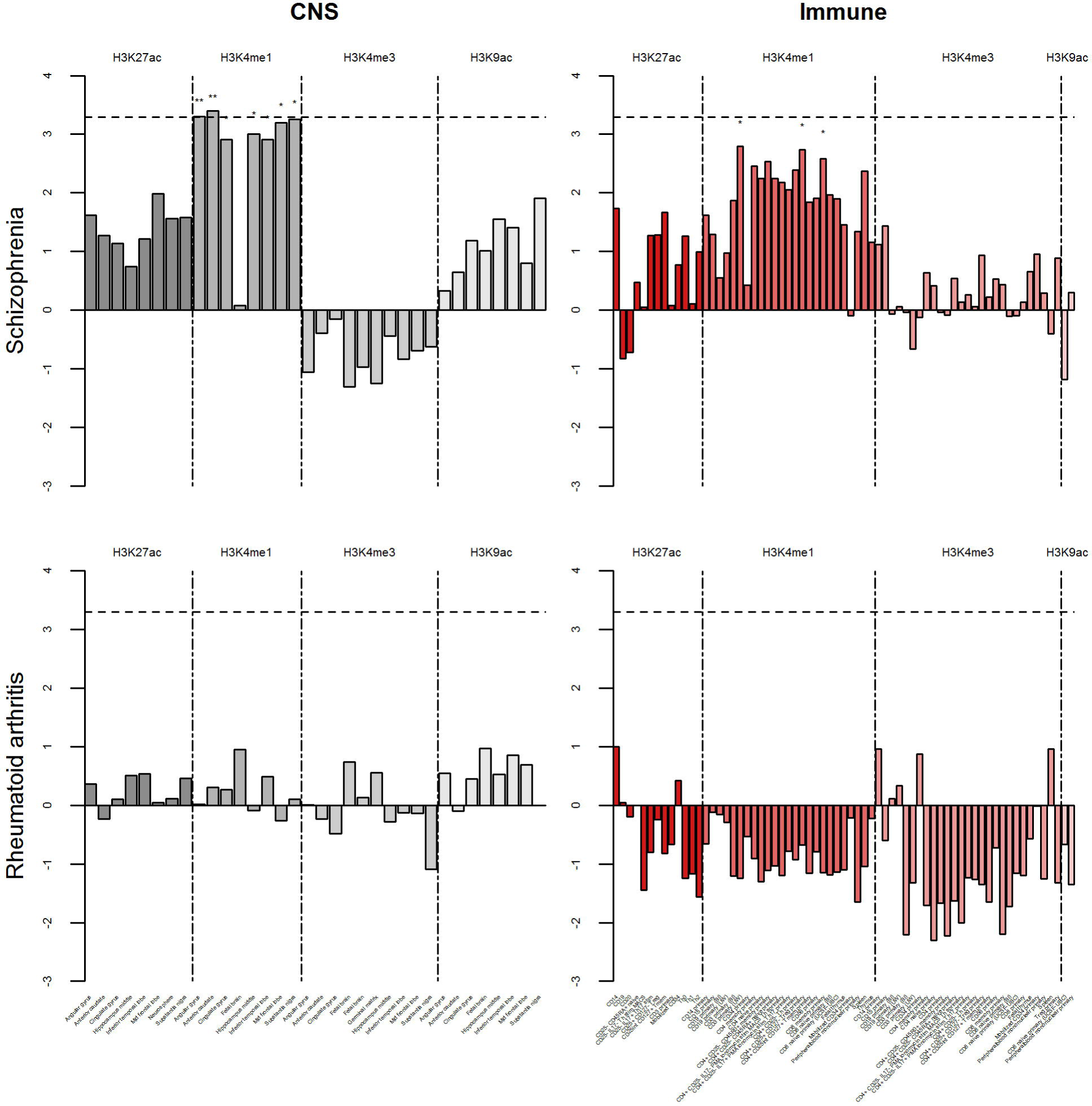
Coefficient Z-score of intersection between union of GTEx eQTLs and cell-type-specific histone marks. Top two graphs show coefficient Z-scores for schizophrenia. Bottom two graphs show the same for rheumatoid arthritis. Grey bars indicate histone marks found in cells from the central nervous system. Red bars represent histone marks found in cells from the immune system. From dark to light, shades of the bars indicate histone marks H3K27ac, H3K4me1, H3K4me3, and H3K9ac. Vertical dotted lines indicate separation between histone marks. One asterisk above the bars indicate annotations passing FDR correction for multiple testing. Two asterisks indicate bars passing Bonferroni correction for multiple testing. Horizontal dotted line indicates Bonferroni threshold for 101 tests.

### Intersection of GTEx eQTLs and tissue-specific differentially expressed genes

We extracted all eQTLs within the top 10% of tissue-specific differentially expressed genes in all 44 GTEx tissues. We then compare the enrichment in GWAS signal for these eQTLs with the genes themselves. The correlation between the coefficients was 0.58 and 0.24 for schizophrenia and rheumatoid arthritis, respectively. For schizophrenia, we see that the eQTL annotation most brain tissues have the highest regression coefficient and Z-score, although none reached the significance threshold (S10 Table). Furthermore, the eQTLs showed larger coefficients compared to the whole genes, although the large standard errors made the difference non-significant. Interestingly, the top 10% differentially expressed genes within the nucleus accumbens showed a significant coefficient comparable to the other brain regions, although the eQTLs for these genes showed a non-significant depletion. For rheumatoid arthritis, whole blood showed the most significant coefficient, however, again failed correction for multiple testing (S11 Figure). Furthermore both the whole genes as the eQTLs for these genes showed a similar regression coefficient.

## Discussion

Stratified Linkage Disequilibrium Score Regression provides a way to partition h^2^_SNP_ into parts explained by (functional) parts of the genome (Finucane et al. 2015). A “full baseline model” containing 24 non-cell-type-specific annotations of SNPs, such as SNPs located in promoters or coding regions, was developed previously for analysis using SLDSR. Here, we added annotations containing eQTLs derived from whole blood and brain tissue into the model, and showed that eQTLs were substantially stronger enriched in their effect on complex traits compared to all categories considered by Finucane *et al* (2015). The complete brain eQTL annotation was significantly enriched in GWAS signal for educational attainment, rheumatoid arthritis, smoking behavior, and schizophrenia. This finding is consistent with previous estimates of eQTL effect enrichment (Davis et al. 2013). Considerable enrichment for eQTLs, even for traits not apparently linked to the brain or immune system (e.g. smoking behavior), suggested that non-trivial eQTL overlap across tissues might be present.

Inclusion of both brain and blood eQTLs into the SLDSR model did not separate the signal into tissue-specific effects. In general, we are not able to clearly identify tissue-specific eQTL signals using these datasets and SLDSR. Our second analysis of eQTL enrichment based on 44 tissue-specific *cis*-eQTL sets, obtained from the GTEx consortium (2015; Aguet et al. 2016), confirms the lack of tissue-specific eQTL enrichment. While an annotation containing all eQTLs identified in GTEx is significantly enriched in its effect on schizophrenia and rheumatoid arthritis (Z=5.501 and Z=3.802, respectively, both *p*<0.001), none of the analyzed brain tissues are enriched beyond all eQTLs in their effect on schizophrenia. Similarly, whole blood eQTLs are not significantly enriched beyond all GTEx eQTLs taken together in their effect on rheumatoid arthritis. Again, these findings are not consistent with the hypothesis of abundant tissue-specific *cis*-eQTLs with effects on complex traits related to the specific tissue in question. Our findings are consistent with a lack of power to detect any tissue-specific eQTL effects. Especially, when contrasted with tissue-specific gene expression levels and tissue-specific histone modifications (Liu et al. 2016; Finucane et al. 2017), tissue-specific eQTLs are of limited value in relating complex traits to a tissue. In fact, considering eQTLs associated with genes expressed in a specific tissue improves our detection of tissue-specific effects. But, while the regression parameter subsets eQTLs for specifically expressed genes have larger effects than the rest of these genes, the significance of the enrichment is weak compared to the significance of the tissue-specific enrichment of the whole gene body plus a 100kb window.

One of the limitations of the study presented here involves the substantial differences in discovery sample size between the tissues, which influences the power to detect eQTLs (Lonsdale et al. 2013). Even within the GTEx tissues, where differences in sample sizes are relatively small compared to eQTLs obtained from Jansen *et al* and Ramasamy *et al,* we still see a significant correlation between the discovery sample size and enrichment of eQTLs in GWAS signal. Several methods have been developed to capitalize on cross-tissue overlap in eQTLs to improve power to detect SNP effects on gene expression within tissue (Flutre et al. 2013; Li et al. 2013). The aim of this paper was to explore the possibilities of assessing the effects of eQTLs expressed in whole blood on presumably brain-related traits, and vice versa. In the analyses with eQTLs for differentially expressed genes, we show that enrichment in GWAS signal is stronger in these eQTLs compared to taking the all SNPs in the same genes. This indicates that eQTLs, irrespective of the tissue in which they were discovered, play an important role in the etiology of complex traits, and do so via the gene they are associated with. This, however, does not take away the need to increase sample sizes when performing tissue-specific discovery of (*cis*-)eQTLs. Tissue specificity, in the end, is a relative judgement best reached based on weighing multiple lines of evidence, among which are differential expression, epigenetic regulation, and eQTLs. For eQTLs to play a large role in determining the tissue-specific effects on complex traits, a continued investment in resources like GTEx is required in order to increase sample sizes for detection, especially in rare tissues.

Our conclusions are limited to *cis*-eQTLs and it is not unlikely that trans-eQTLs behave differently in terms of tissue-specificity. We do find evidence for possible enrichment for eQTLs that intersect with tissue-specific H3K4me1 histone marks in the brain, but also immune cells, in their effect on schizophrenia but not rheumatoid arthritis. This means that eQTLs in H3k4mel marks are enriched in their effect on schizophrenia above the expected enrichment based on the fact that these SNPs are both eQTLs and located in H3K4me1 histone marks. What is of substantial interest is that the enrichment in GWAS signal appears specific to H3K4me1 marks, and no other Histone marks, suggesting that these marks specifically can aid in prioritizing genomic regions in which tissue-specific eQTLs may reside. Though, again, the totality of evidence is inconclusive on the relevance of tissue-specific eQTLs to variation in complex traits.

Our results are consistent with, and complimentary to, a study investigating the genetic correlation between gene expression levels across 15 tissues (Liu et al. 2016). This study revealed substantial correlations between *cis*-genetic effects on gene expression across 15 tissues (Liu et al. 2016). Our analyses confirmed the value of using whole blood as discovery tissue for detection of c/s-eQTLs and further demonstrated the usefulness of techniques that use c/s-eQTLs discovered in whole blood to study the etiology of complex traits related to different tissues (Gamazon et al. 2015; Gusev et al. 2016). The results presented here highlight the overlap of *cis*-eQTL effects across tissues on a genome-wide level. However, the effect of a c/s-eQTL might vary substantially across tissues for individual genes (Grundberg et al. 2012). Our conclusions are based on genome-wide enrichments and therefore should not be interpreted as limited evidence for tissue-specific eQTL effects for individual genes. Therefore, eQTL discovery in the tissue most relevant to a specific trait or disorder remains important to further our understanding of the genetic regulation of tissue-specific gene expression. What is also clear is that to discover those tissue-specific eQTLs that are of relevance to the interpretation of GWASs of complex traits, tissue-specific eQTL discovery needs to be refined.

The practice of, as a post-hoc analysis to GWAS, performing eQTL lookup in a specific tissue linked to a trait, when larger dataset for other accessible tissues are available, may be suboptimal. In fact, one may prefer to perform a lookup in the overlap between histone modifications in a relevant tissue and eQTLs regardless of tissue. One can further consider utilizing eQTLs to link GWAS findings to a gene, and subsequently consider the differential expression of a gene to identify the tissue in which the gene is most likely to act in effecting the trait. Tissue-specific differential gene expression vastly outperforms eQTLs in tagging regions of the genome enriched in their effect on complex traits (Finucane et al. 2017).

It is also evident that a limited dichotomous definition of eQTL/no-eQTL may be insufficient to identify tissue-specific eQTL effects. An evident improvement would be to compute the *difference* in eQTL effect on expression of the gene between tissues, and perform inference based on this difference in effect. eQTLs are strongly enriched SNPs, with clear biological function and utility for the translation of GWAS findings, though tissue-specific eQTL mechanisms remain elusive. The discovery of tissue-specific eQTL effects, which can aid in linking complex trait to tissue, may require novel research strategies.

## Supplemental Data

Supplemental Data includes 4 figures and 7 tables

## Compliance with Ethical Standards

### Funding

MGN is supported by the Royal Netherlands Academy of Science Professor Award (PAH/6635) to DIB. HFI is supported by the “Aggression in Children: Unraveling gene-environment interplay to inform Treatment and InterventiON strategies”(ACTION) project. ACTION receives funding from the European Union Seventh Framework Program (FP7/2007–2013) under grant agreement no 602768. MB is supported by a University Research Chair of the Vrije Universiteit. The discovery of blood eQTL was funded by the US National Institute of Mental Health (RC2 MH089951, principal Investigator PFS) as part of the American Recovery and Reinvestment Act of 2009. We thank T. Lehner (National Institute of Mental Health) for his support. We acknowledge Hillary Finucane, Raymond Walters and Benjamin Neale for critical comments on our methods, design and manuscript.

### Ethical approval

This article does not contain any studies with human participants or animals performed by any of the authors.

### Web Resources

Age at menarche summary statistics, https://www.reprogen.org/data_download.html

Blood eQTLs, https://eqtl.onderzoek.io/

Brain eQTLs, http://www.braineac.org/

Coronary artery disease summary statistics, https://www.cardiogramplusc4d.org/data-downloads/

Crohn’s disease and ulcerative colitis summary statistics, https://www.ibdgenetics.org/downloads.html

Educational attainment summary statistics, http://www.thessgac.org/data

Full baseline model LD scores, http://data.broadinstitute.org/alkesgroup/LDSCORE/

GTEx dataset, http://www.gtexportal.org/home/datasets

Height and BMI summary statistics, https://www.broadinstitute.org/collaboration/giant/index.php/GIANT_consortium_data_files

LDL levels summary statistics, https://www.broadinstitute.org/mpg/pubs/lipids2010/

Rheumatoid arthritis summary statistics, http://plaza.umin.ac.jp/yokada/datasource/software.htm

Schizophrenia and smoking behavior summary statistics, https://www.med.unc.edu/pgc/results-and-downloads

SLDSR software, https://github.com/bulik/ldsc/

